# Correspondence between resting state activity and brain gene expression

**DOI:** 10.1101/021071

**Authors:** Guang-Zhong Wang, T. Grant Belgard, Deng Mao, Leslie Chen, Stefano Berto, Todd M. Preuss, Hanzhang Lu, Daniel H. Geschwind, Genevieve Konopka

## Abstract

The relationship between functional brain activity and gene expression has not been fully explored in the human brain. Here, we identify significant correlations between gene expression in the brain and functional activity by comparing fractional Amplitude of Low Frequency Fluctuations (fALFF) from two independent human fMRI resting state datasets to regional cortical gene expression from a newly generated RNA-seq dataset and two additional gene expression datasets to obtain robust and reproducible correlations. We find significantly more genes correlated with fALFF than expected by chance, and identify specific genes correlated with the imaging signals in multiple expression datasets in the default mode network. Together, these data support a population-level relationship between regional steady state brain gene expression and resting state brain activity.

**HIGHLIGHTS:**

- Gene expression is correlated with resting state activity in human brain.
- Specific genes are correlated with activity in the default mode network.
- Activity-correlated genes are enriched in neurons.
- Regionally-patterned gene expression is enriched for activity-correlated genes.

## INTRODUCTION

Understanding the molecular underpinnings of complex human brain circuits requires integration of multiple forms of high-dimensional data, ranging from functional brain imaging(Medland et al., 2014; Scott-Van Zeeland et al., 2010) to genetic and genomic analyses. Seminal work in neuroimaging over the last decade has revealed that human brain activity can be defined by its network properties (Buckner and Krienen, 2013; Goni et al., 2014; Power et al., 2011; Sporns, 2013). Differences in gene regulation and expression have also been related to brain function in many species (Barchuk et al., 2007; Chandrasekaran et al., 2011; O’Connell and Hofmann, 2011) and to the evolution of human higher cognitive functions (Geschwind and Konopka, 2012; Geschwind and Rakic, 2013; Konopka et al., 2012; Preuss, 2012; Somel et al., 2013; Wang and Konopka, 2013). Here, we link these diverse levels of analysis by asking whether gene expression in human neocortex is correlated to a property of functional networks defined by fMRI in humans (Figure 1A).

**Figure 1.**
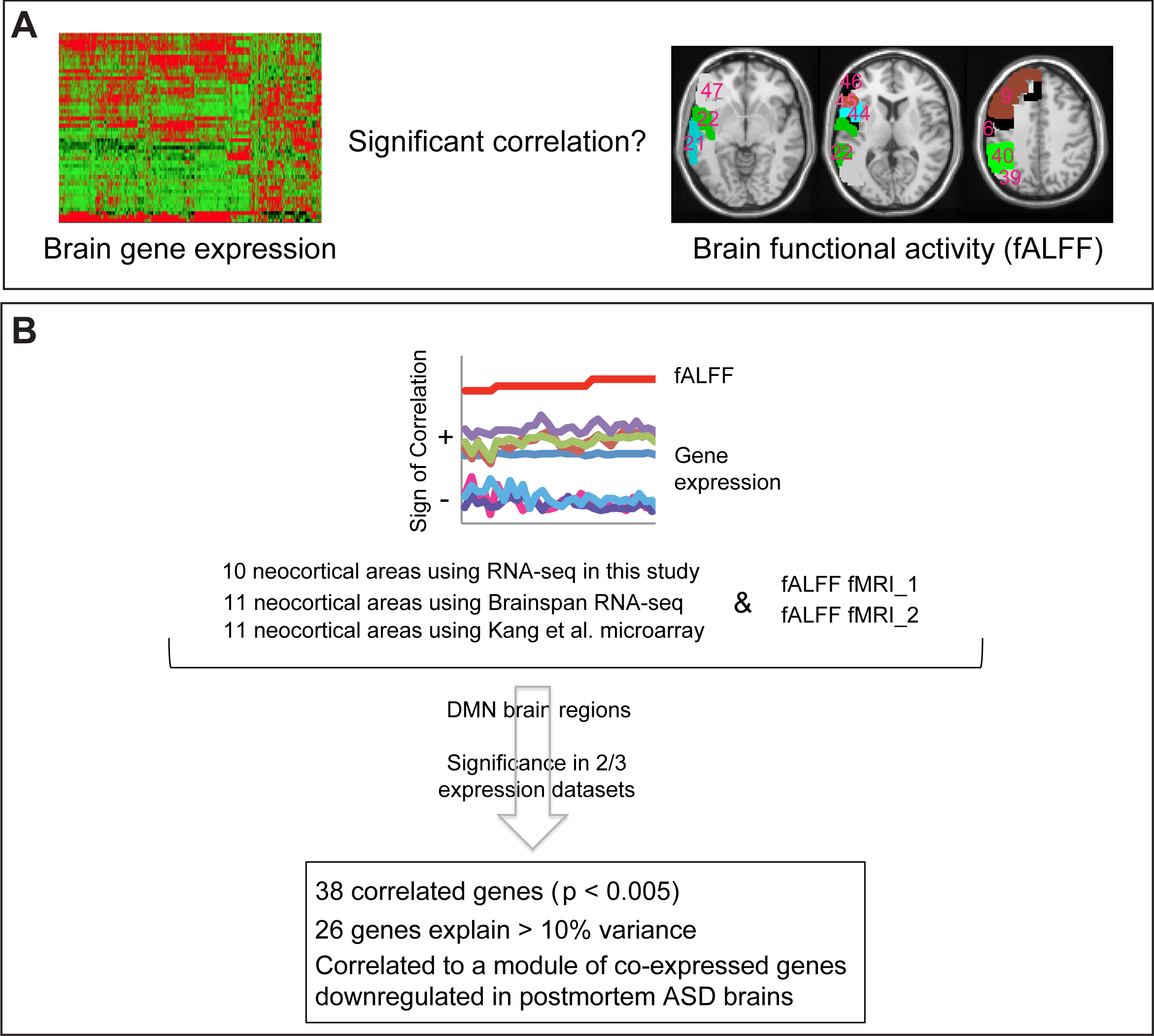
Schematic of data analysis and results. (A) Significant correlations were investigated between gene expression and fMRI signal (fALFF) among different brain regions. Brodmann areas from our RNA-seq dataset are indicated in the fALFF schematic. (B) Three gene expression studies were used. The RNA-seq data in this study contained 52 left hemisphere samples in total from 10 neocortical regions, including 25 samples from 5 DMN regions; the Brainspan RNA-seq data contained 129 left hemisphere samples from 11 neocortical regions, including 72 samples from 6 DMN regions; and the Kang et al. microarray data contained 378 samples from 11 neocortical regions in both left and right hemispheres, including 212 samples from 6 DMN regions. Two resting state fMRI datasets were used, containing 198 and 84 individuals, respectively. Significantly correlated genes were identified using a Fisher r-to-z transformation, p < 0.005. Enrichment or significance was determined based on consistency in two out of three expression datasets.

Because gene expression studies in the human brain are limited to post-mortem tissue, it might be difficult to meaningfully relate brain gene expression to brain activity measured during specific task-based paradigms. We therefore chose only to include brain activity measurements obtained during the resting state when individuals are not being asked to carry out a specific task. In addition, we primarily focus on a subset of cortical regions active in one particular network that is active during a resting state, the default mode network or DMN. The DMN is one of several brain networks active during a resting state that may indicate cognitive function that is either non-directed or self-referential (Buckner et al., 2008). Previous work has indicated that genetic factors might regulate DMN function and disruption to DMN function has been observed in highly heritable brain disorders such as autism (Washington et al., 2014).

We used specific activity values (fractional Amplitude of Low Frequency Fluctuations or fALFF, the fraction of total amplitude in the 0.01-0.08 Hz frequency band) obtained during resting state fMRI as our proxy for brain activity. Previous work has indicated that the blood-oxygen-level dependent contrast, or BOLD signal, measured by fMRI is a direct reflection of neuronal activity (Amit and Romani, 2007). However, it is unknown which genes, if any, might be orchestrating the neuronal activity measured by this fMRI signal and whether these gene expression patterns might be directly attributable to neuronal gene expression patterns rather than secondary ones in other cells types. We hypothesized that neuronal patterns of gene expression are driving cellular patterns of activity that can be observed during brain imaging. Here, we identify the specific genes correlated with brain activity in brain regions that are part of the DMN. We find that these correlated genes are more likely to be enriched in neurons relative to other major CNS cell types such as endothelial cells that could also be involved in regulating BOLD signal. We also observe that the correlated genes have previously been reported to have altered expression in autism. Thus, by focusing our study specifically on the DMN and using multiple sources of brain gene expression and imaging signals, we have identified a core set of genes that might not only be important for the brain activity in the DMN at rest but might also be preferentially affected in cognitive disorders such as autism.

## RESULTS

### Expression of individual genes correlate with fMRI signals

To assess the relationship between gene expression regulation and human brain activity, we first generated a human neocortical RNA-sequencing (RNA-seq) dataset. We carried out RNA-seq of ∼50 adult human postmortem neocortical brain samples representing 10 neocortical brain regions: pre-motor cortex (PMV; BA6 (Brodmann Area 6)), dorsolateral prefrontal cortex (DLPFC; BA9), posterior middle temporal gyrus (pMTG; BA21), posterior superior temporal gyrus (pSTG; BA22), angular gyrus (AG; BA39), supramarginal gyrus (SMG; BA40), pars opercularis (POP; BA44), pars triangularis (PTr; BA45), middle frontal gyrus (MFG; BA46), and pars orbitalis (POrB; BA47). For each brain region, three or more samples from left adult brain hemispheres were collected (ages range from 33 to 49) (Tables S1 and S2). An example of the gene expression profiles across our RNA-seq samples is illustrated in Figure S1. In addition to our own RNA-seq samples from 10 neocortical areas (5 of which have DMN involvement), we used two additional large scale transcriptome datasets in this analysis: RNA-seq data from Brainspan (www.brainspan.org) and microarray data from Kang et al. (Kang et al., 2011), which each consist of 11 neocortical areas (6 with DMN involvement) (Figure 1B).

We next compared these data to resting state fMRI data (Biswal et al., 2010), computing for every gene the correlation between its expression level across cortical areas and fALFF in those areas (Zou et al., 2008), which is considered a proxy for regional brain activity, from two large publicly available fMRI datasets (herein referred to as fMRI_1 and fMRI_2) (Tables S3 and S4; Figure 1B) (Biswal et al., 2010). The mean fALFF from all individuals was computed for each cortical area separately from each of these two fMRI datasets. We then computed Pearson correlations between these values and the expression levels of each gene across the relevant cortical areas in each of three gene expression datasets and two imaging datasets. Although the samples used in Brainspan are partially a subset of the samples used in Kang et al., there was great similarity between the gene expression-fALFF correlations in our RNA-seq data and Brainspan (Pearson’s rho = 0.87) and between the gene expression-fALFF correlations in our RNA-seq data and the Kang et al. data (Pearson’s rho = 0.90) for the 38 genes reported here. Hence, the lack of complete independence of Brainspan and Kang et al. does not substantially affect this analysis.

Only genes having significant correlations to fALFF (Fisher r-to-z transformation, two-tailed p < 0.005) in at least two of the three gene expression datasets for both imaging datasets were included for further analysis. Additionally, we used directionality as an evaluation of the consistency of our method as we expect that the positive and negative correlations should be consistent for the same genes across datasets.

Using all brain regions available, we found that 126 genes met our stringent criteria of replication across 2/3 brain expression datasets and 2 fMRI studies; however, only 11 genes were consistent with the additional requirement of direction (positive or negative) across all six pairs of datasets (three gene expression by two fALFF vectors). This was not surprising as this large number of brain regions was sequenced based on tissue availability rather than on functional connectivity. We therefore focused our comparisons on the brain regions with known involvement in the DMN, which should primarily be active during a resting state, the condition the fMRI data are derived from. While other resting state networks could be evaluated, we chose the DMN due to the availability of gene expression in specific brain regions from multiple datasets including our newly generated dataset.

Using the DMN regions, we found more genes than expected by chance to be significantly correlated with fALFF in each of the three datasets (permutation tests of fALFF assignments; empirical p = 0.04 for our RNA-seq, p = 0.004 for Kang et al. and p = 0.03 for Brainspan, Figure 2A-C). We carried out permutation experiments (1000 times) and observed that fewer than 4 genes (3.2 genes) would be expected by chance to have such a correlation using these criteria as we found using all brain regions. Using the DMN regions, we identified 38 fALFF-correlated genes using these criteria (p < 0.01, empirical p-value from permutation described above, Table S5.) Strikingly, all of these 38 genes are correlated in the same direction in both fMRI datasets, with 12 of them positively correlated and 26 of them negatively correlated with fALFF. Additionally, these 38 genes show significantly higher correlation with gene expression in the brain regions examined than in non-nervous system tissue samples (p < 0.01 for fMRI_1 and fMRI_2, respectively), and none of them are known human housekeeping genes (Eisenberg and Levanon, 2013). Moreover, sex of the donor did not substantially affect correlations between gene expression and fMRI activity (Figure S2). The gene-fALFF correlations are highly similar in the two fMRI datasets (Pearson’s rho > 0.99 on average comparing gene-fALFF correlations in one fMRI dataset to the other, Figure 3), indicating the robustness of this analysis across imaging cohorts. Remarkably, the expression pattern of 26 of the 38 genes can each explain 10% or more of the statistical variance in fALFF (Table 1).

**Table 1.** The 26 genes with expression correlated to fALFF in the same direction across datasets and >10% variation explained on average per gene.

| Gene Name | Sign | Variance explained ($R^2$) fMRI_1 | | | Variance explained ($R^2$) fMRI_2 | | | DE gene |
| --- | --- | --- | --- | --- | --- | --- | --- | --- |
|  |  | In this study | Brainspan | Kang et al. | In this study | Brainspan | Kang et al. |  |
| *CCDC41* | - | 0.42 | 0.14 | 0.08 | 0.53 | 0.13 | 0.09 |  |
| *NPTXR* | - | 0.30 | 0.06 | 0.07 | 0.56 | 0.04 | 0.06 |  |
| *TP53I11* | - | 0.29 | 0.10 | 0.11 | 0.49 | 0.08 | 0.10 | Yes |
| *PIM1* | + | 0.26 | 0.14 | 0.07 | 0.13 | 0.14 | 0.08 |  |
| *DRD2* | - | 0.22 | 0.15 | 0.20 | 0.31 | 0.10 | 0.19 | Yes |
| *GLRA2* | - | 0.21 | 0.28 | 0.12 | 0.43 | 0.23 | 0.10 |  |
| *NR2F2* | - | 0.20 | 0.38 | 0.24 | 0.26 | 0.35 | 0.23 | Yes |
| *NECAB2* | - | 0.19 | 0.16 | 0.12 | 0.36 | 0.11 | 0.11 | Yes |
| *PEA15* | - | 0.17 | 0.17 | 0.05 | 0.30 | 0.16 | 0.04 |  |
| *YPEL1* | - | 0.17 | 0.12 | 0.08 | 0.35 | 0.09 | 0.06 |  |
| *EPB41L4B* | - | 0.16 | 0.13 | 0.06 | 0.31 | 0.12 | 0.07 |  |
| *FNBP1L* | - | 0.14 | 0.16 | 0.06 | 0.30 | 0.15 | 0.05 |  |
| *STAC2* | + | 0.14 | 0.11 | 0.15 | 0.26 | 0.13 | 0.15 | Yes |
| *HTR2C* | - | 0.11 | 0.14 | 0.11 | 0.23 | 0.11 | 0.07 | Yes |
| *TNNT1* | - | 0.11 | 0.16 | 0.08 | 0.17 | 0.13 | 0.07 |  |
| *CACHD1* | - | 0.10 | 0.12 | 0.09 | 0.24 | 0.13 | 0.08 |  |
| *NEFH* | + | 0.10 | 0.15 | 0.10 | 0.19 | 0.18 | 0.09 | Yes |
| *SYT2* | + | 0.10 | 0.14 | 0.19 | 0.25 | 0.19 | 0.20 | Yes |
| *KCNS1* | + | 0.09 | 0.15 | 0.12 | 0.20 | 0.15 | 0.10 | Yes |
| *SCN1B* | + | 0.08 | 0.12 | 0.11 | 0.21 | 0.15 | 0.10 | Yes |
| *SPC25* | - | 0.08 | 0.25 | 0.04 | 0.16 | 0.23 | 0.04 |  |
| *RAB37* | + | 0.07 | 0.12 | 0.15 | 0.15 | 0.13 | 0.17 |  |
| *RAPGEF4* | - | 0.07 | 0.21 | 0.07 | 0.13 | 0.20 | 0.06 |  |
| *KCTD4* | - | 0.06 | 0.18 | 0.19 | 0.19 | 0.16 | 0.17 | Yes |
| *ARHGAP6* | - | 0.04 | 0.27 | 0.09 | 0.14 | 0.22 | 0.06 |  |
| *GPC4* | - | 0.04 | 0.22 | 0.09 | 0.14 | 0.21 | 0.08 |  |

**Figure 2.**
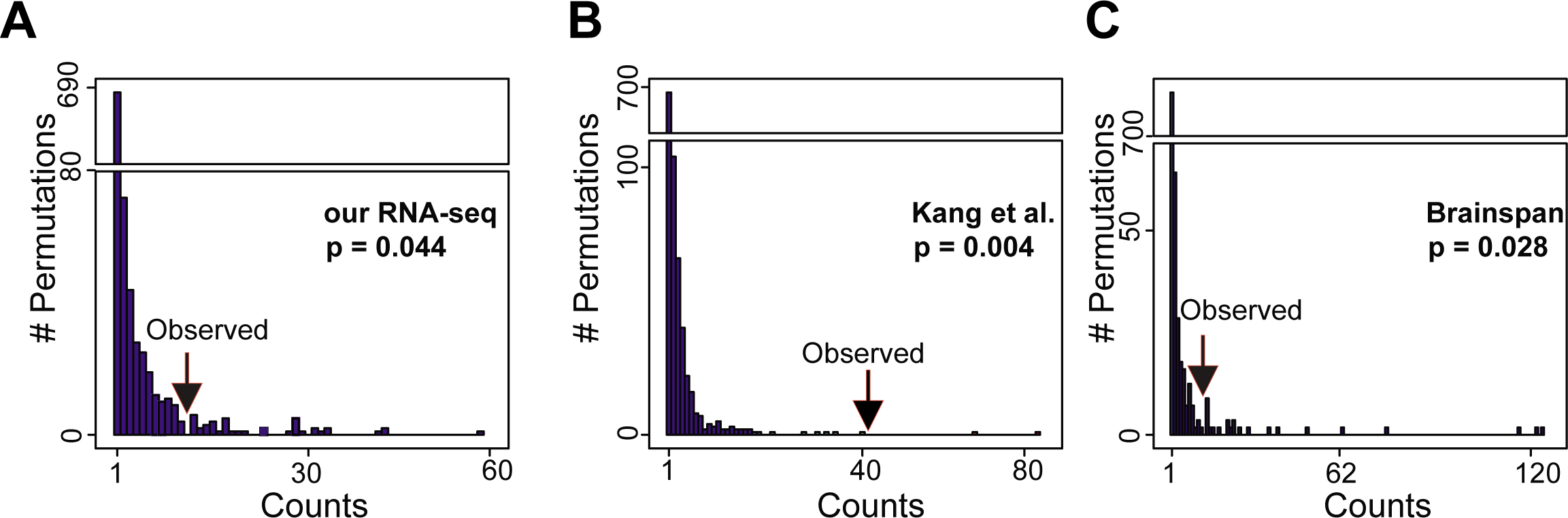
Significantly more genes than expected are correlated in expression to fALFF. This indicates that this particular patterning of genes, whose expression tracks regional brain activation as measured by fALFF, is much more common than random patterns. The arrow indicates the observed number of correlated genes in each dataset ((A) our RNA-seq, (B) Kang et al., and (C) Brainspan). Bar plots indicate the number of correlated genes in the randomized dataset. Fold enrichment is 5, 25 and 6 for our RNA-seq, Kang et al. and Brainspan respectively. 1,000 permutations were carried out and the empirical significance level is indicated.

**Figure 3.**
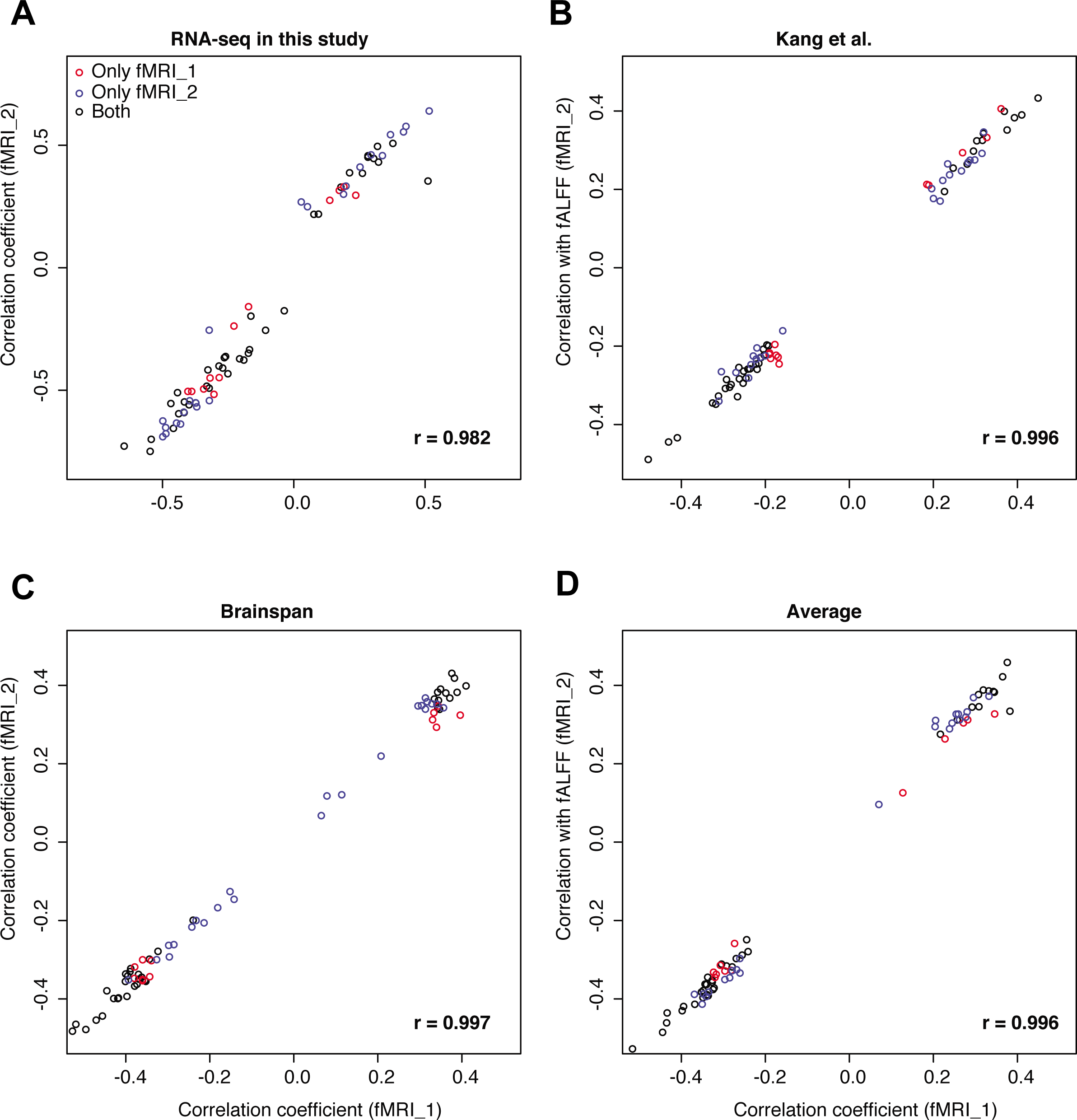
Replication of fALFF-correlated genes using two fMRI datasets. Pearson correlations between fMRI datasets using (A) our RNA-seq data, (B) Kang et al., (C) Brainspan and (D) an average of all three expression datasets. Red points indicate fALFF-correlated genes in only the fMRI_1 dataset. p < 1e-10 for all the correlations. Blue points indicate fALFF-correlated genes in only the fMRI_2 dataset. Black points indicate fALFF-correlated genes in both fMRI_1 and fMRI_2. These data demonstrate that even when correlations reach significance in only one of the two fMRI datasets, the correlation levels are nonetheless highly similar.

Several of these fALFF-correlated genes play roles in brain function and disease. In Figure 4A, we illustrate a few of the most highly positive and negatively correlated genes. Many of these genes have previously been implicated in neuronal activity or dysfunction, which is consistent with some of these genes possibly having a role in resting state activity that is disrupted in cognitive diseases. *SYT2* encodes an integral membrane protein of synapses and can increase the dynamic range of synapses by maximizing calcium-evoked neurotransmitter release (Kochubey and Schneggenburger, 2011). *SCN1B*, which regulates the Na+ current in neurons, has been implicated in epilepsy and Dravet syndrome (Carranza Rojo et al., 2011; Fujiwara et al., 2003; Marini et al., 2011; Wallace et al., 1998). *GLRA2* is a ligand-gated ion channel that is widely expressed in the central nervous system and plays an important role in the inhibition of neuronal firing (Young-Pearse et al., 2006). *DRD2* as well as *HAPLN4* (see Table S5) were recently identified in the largest schizophrenia GWAS to date as two of the genes near the highly associated loci (Schizophrenia Working Group of the Psychiatric Genomics Consortium. 2014), however with so few fALFF-correlated genes we were underpowered to systematically assess this relationship.

**Figure 4.**
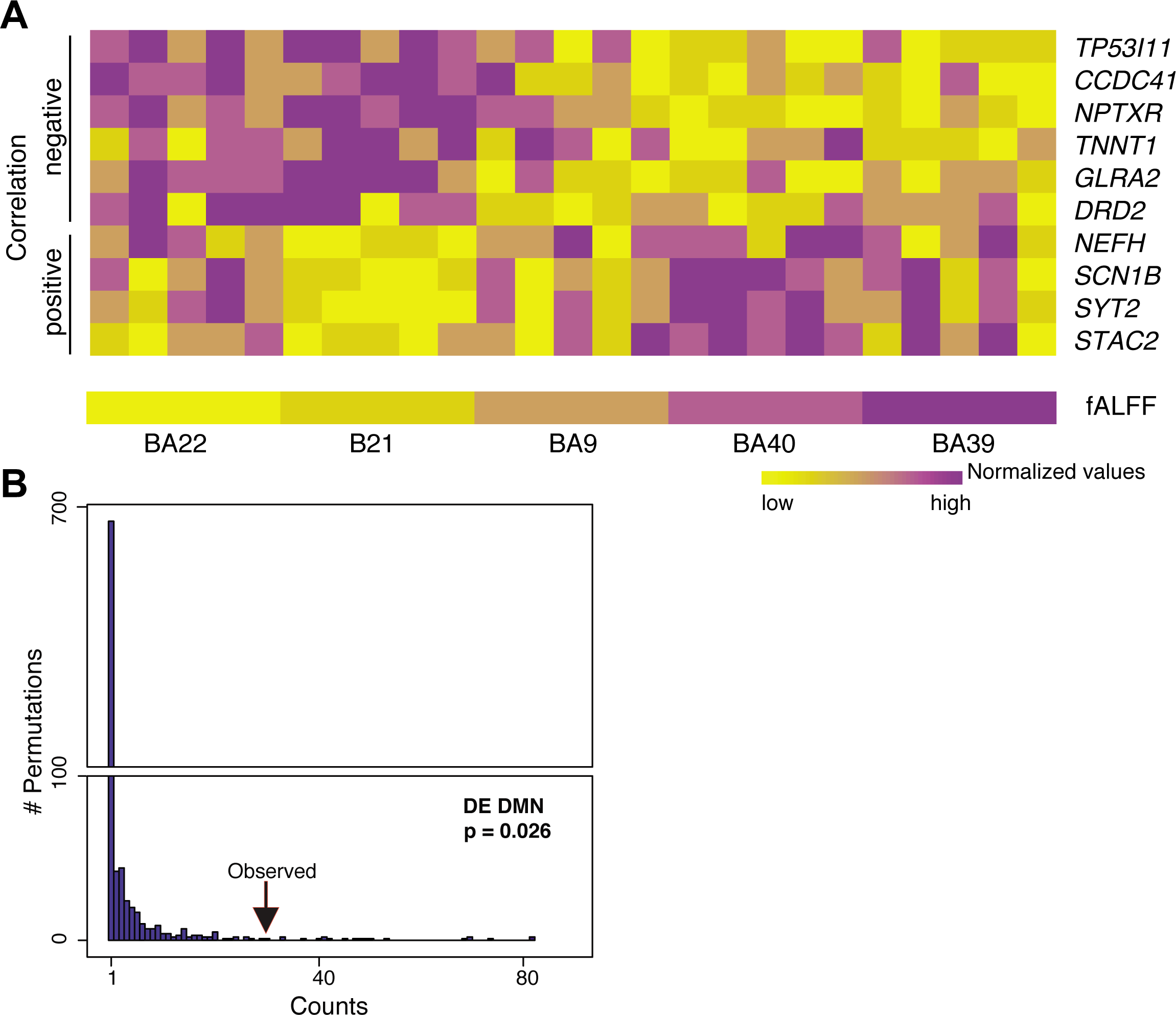
Expression of individual genes is correlated with fALFF signals and these correlations are greater for regional DE genes. (A) Heatmap view of some of the top fALFF-correlated genes. Yellow indicates smaller values in the fALFF data or gene expression and purple indicates higher values. *TP53I11*, *CCDC41, NPTXR, TNNT1, GLRA2,* and *DRD2* are negatively correlated with fALFF and *NEFH, SCN1B*, *SYT2,* and *STAC2* are positively correlated with fALFF. fALFF values from fMRI_1 are presented and the order of the normalized fALFF values is the same in both fMRI datasets. (B) Regional DE genes are more significantly correlated with fALFF than expected. Bar plots indicate DE genes correlated with fMRI in our RNA-seq data set. The arrow indicates the observed number of correlated DE genes. Bar plots indicate the number of correlated genes in the randomized dataset. The fold enrichment is 8.5 for DMN regional DE genes. 1,000 permutations were carried out and the empirical significance level is indicated.

### fALFF-correlated genes have greater expression in neurons

Based on these connections between the most highly correlated genes and neuronal function, we examined a genome-wide database of brain cell type specificity (Zhang et al., 2014). For the 34 genes having available cell-type expression data out of these 38 fALFF-correlated genes, we determined the cellular preference of expression for each gene based on the highest RPKM value among the seven cell types in Zhang et al (Zhang et al., 2014). We find that a plurality (14) of the highly fALFF-correlated genes are preferentially expressed in neurons (Figure S3 and Table S6, p = 0.003; Fisher’s exact test) and >85% (12 out of the 14 genes with preferential neuronal expression) of these genes have at least two times higher expression in neurons compared to the maximum expression of other cell types. As only two genes, *TNNT1* and *GPD2* (Table S6), are most highly expressed in endothelial cells and fALFF correlated genes are depleted, as a group, in endothelial cells (p = 0.03; two-tailed Fisher’s exact test), we conclude that these correlations between fALFF and gene expression are not likely driven by vascular density, but rather by a signature derived from neuronal function. This is consistent with reports that fALFF is a specific measure of neuronal activity (Zou et al., 2008).

### fALFF-correlated genes overlap genes downregulated in ASD

Since previous work has shown that the DMN is disrupted in cognitive disorders such as autism spectrum disorders (ASD) (Jung et al., 2014), we examined whether salient ASD cortical gene expression patterns overlap with the highly fALFF-correlated genes. We find 9 fALFF-correlated genes that are contained within the autism gene co-expression module, asdM12, a set of genes downregulated in autism previously identified from postmortem ASD brain tissue (Table S7, p = 8e-5; Fisher’s exact test) (Voineagu et al., 2011). All but one of these genes are negatively correlated with resting state fALFF (Table S7). By definition, the genes within the asdM12 module are highly co-expressed with one another across the temporal and frontal cortex of multiple individuals. We therefore computed the expression correlations of the 38 fALFF-correlated genes. By permuting pairs of genes, we find that the fALFF-correlated genes show approximately 2 fold higher co-expression than other genes (p < 1e-3) and there is no bias towards positive or negative correlations (Figure 4A, Figure S4 and S5). As the genes in the asdM12 module are enriched with neuronal and synaptic markers and tend to be downregulated in autism brains(Voineagu et al., 2011), our results again highlight the importance of these correlated genes in normal brain function. For example, *RAPGEF4* (also known as *EPAC2*) is one of the fALFF-correlated genes that is contained in the asdM12 module. Rare variants of *RAPGEF4* have also been reported in patients in autism (Bacchelli et al., 2003). The function of *RAPGEF4* is to serve as a cAMP-dependent guanine nucleotide exchange factor and dysfunction of *RAPGEF4* is associated with spine remodeling and synapse function (Kawasaki et al., 1998; Penzes et al., 2011). Therefore, *RAPGEF4* plays an important role in brain function that is further emphasized by correlation of its expression to resting state brain activity.

### fALFF-correlated genes are enriched in brain regional DE genes

Previous human brain transcriptome studies have noted a dearth of genes differentially expressed between cortical areas (Jaffe et al., 2015; Pletikos et al., 2014), and much of the differential expression driving cortical diversity likely derives from laminar specificity. However, there has been very little functional characterization of these cortical patterned genes or ideas put forth as to how their expression pattern may contribute to overall brain function. Based on our approach of correlating fALFF and gene expression, we have identified differential gene expression signatures between cortical areas that correspond to the differential brain activity. To confirm this, we assessed pairwise brain regional differential expression in these DMN-involved cortical areas (Table S8). We identified 545 genes differentially expressed between these areas. We find that 29% of the identified fALFF-correlated genes are indeed differentially expressed (DE) genes (11/38, p = 6e-14. Twenty of the fALFF-correlated genes are also differentially expressed during neocortical developmental (Table S5) (Jaffe et al., 2015). We find that these patterned genes have significantly higher correlations with fALFF than non-patterned genes (Figure S6, p < 1e-10 in all the comparisons). Additionally, there are more genes than expected that are correlated with fALFF signals among the differentially expressed genes (Figure 4B, p < 0.05 in permutation experiments). Together, these results are consistent with patterned genes having a functional role in steady state brain activity.

## DISCUSSION

This is one of the first studies to directly correlate the transcriptome to measurements of human brain activity. Here, we correlate RPKM, a measurement of steady state gene expression with fALFF, a measurement for resting state activity within a single brain region and find a specific set of highly-correlated genes within the DMN. A recent study has also found correlation between cortical gene expression and two functionally defined brain networks (Cioli et al., 2014). In contrast to our study, this other study finds that two of our top positively correlated genes with the DMN, *SCN1B* and *SYT2*, are correlated with what these authors define as a functional visual-sensorimotor-auditory network. Therefore, while this other study did not use fMRI values correlated to gene expression as we have done, these findings complement our work by identifying some of the same genes corresponding to known functionally connected networks based on brain imaging. Finally, another recently published study used a brain connectivity dataset derived from resting state fMRI to identify ∼78 genes highly corrected with multiple brain resting state networks (Richiardi et al., 2015). We find that 4 of the genes they identify overlap with our set of 38 genes that can explain >10% variation of fALFF (p =1.65e-5; Fisher’s exact test, *NECAB2, NEFH, SCN1B,* and *SYT2*). Three of these genes are positively correlated with fALFF (*NEFH, SCN1B,* and *SYT2*) and one is negatively correlated with fALFF (*NECAB2*). The overlap of these 4 genes supports the robustness of our study and is validation of our findings despite the different methods and brain regions used in the other study. Moreover, since *SCN1B* and *SYT2* were found in all three studies, these data suggest that functional assessment of the function of these specific genes relevant to resting state brain function is highly warranted.

In contrast to the functional connectivity approaches used in the other two studies, our use of fALFF allows for direct measurement of the amplitude of brain activity within a single brain region, rather than between-region correlations (Zou et al., 2008). In addition, we use three independent human brain gene expression datasets (two of which utilize RNA-seq) to provide replication, and two large-scale independent fMRI datasets. A limitation of all of these studies is that one cannot directly address whether resting state brain activity influences gene expression or *vice versa* given the use of post-mortem brain tissue for expression. However, the identification of these genes whose expression is related to brain networks opens the possibility of exploring such directional functional relationships between gene expression and brain activity in animal models.

Our focus on the DMN rather than on several functional networks allows us to prioritize a specific tractable set of genes for the future study of DMN function and its disruption in disease. Because brain regions were defined by Brodmann areas in one of our compared gene expression datasets and gyral landmarks in the other two datasets, we cannot directly compare these brain regions. However, we found remarkable consistency of the fALFF correlated genes across all three gene expression datasets and did not find greater correspondence between the two datasets using gyral landmarks. Such robustness of our findings underscores the stringency of our approach and the utility of focusing on one brain network (the DMN in this case) with multiple gene expression datasets that overlap with the network but not necessarily with one another. Moreover, due to limited availability of additional brain tissue from independent studies for gene expression that correspond to additional DMN regions, future studies will need to explore the robustness of our results throughout the DMN and other resting state networks. However, as the regions of interest among gene expression studies in fact capture different parts of the DMN (Tables S3 and S4), there may be some overlap in gene correlations in additional areas. Finally, this study provides a framework for future investigations into cognitive disorders with altered DMN function such as ASD. The future identification of correlations between fMRI measures and gene expression in ASD brains will determine whether the correlated genes we identified in this study have altered patterns in a disease state.

### Experimental Procedures

#### Sample Information

Post-mortem tissue from ten neocortical brain regions from adult males was used in this study. These regions are pre-motor cortex (PMV; BA6 (Brodmann Area 6)), dorsolateral prefrontal cortex (DLPFC; BA9), posterior middle temporal gyrus (pMTG; BA21), posterior superior temporal gyrus (pSTG; BA22), angular gyrus (AG; BA39), supramarginal gyrus (SMG; BA40), pars opercularis (POP; BA44), pars triangularis (PTr; BA45), middle frontal gyrus (MFG; BA46), and pars orbitalis (POrB; BA47). Detailed sample information is included in Table S1.

#### Sample preparation and RNA sequencing

Frozen tissue samples from human post-mortem brains were used. Samples were dissected from frozen tissue pieces provided by the Maryland Brain and Tissue Bank. Total RNA was extracted using QIAGEN miRNeasy kits according to the manufacturer’s instructions. All RNA samples were examined for quantity and quality by NanoDrop and Bioanalyzer (Agilent). RINs (RNA integrity numbers) are included in Table S1. 1 μg of total RNA from each sample was used, and samples were randomized for RNA extractions and sequencing. Libraries were prepared using an Illumina TruSeq kit according to the manufacturer’s instructions. Samples were barcoded and 3 barcoded samples were run per lane on an Illumina HiSeq according to the manufacturer’s instructions. Single-end 100bp reads were aligned in a staged manner. First, reads were aligned with tophat2 using default options to a reference sequence comprised of mitochondrial DNA, mitochondrial cDNA, and ribosomal cDNA (GRCh37, Ensembl release 67). These reads were excluded from further analysis. The reads that did not map in the first iterations were then mapped using tophat2 to GRCh37 using the GTF provided in Ensembl release 67. RPKMs were then quantified using cufflinks, providing gene definitions (Ensembl release 67 GTF), and enabling several options (fragment bias correction, multi-read correction, upper quartile normalization, and compatible hits normalization). Gene-level RPKM quantifications were used for all downstream processing. Scaled by multiplying the appropriate factor to set each 80^t^^h^ percentile RPKM to be the geometric mean 80^th^ percentile RPKM among samples. The NCBI Gene Expression Omnibus (GEO) accession number for the RNA-seq data reported in this manuscript is GSE58604. Since correlative measures are highly sensitive to zero values, we used the 17,469 genes that were expressed at a nonzero level in each sample. We repeated analyses with and without outlier removal and found it had no substantial impact on the results with these data.

#### fMRI data processing and analysis

The fMRI data were downloaded from The 1000 Functional Connectomes Project (Biswal et al., 2010) (http://www.nitrc.org/frs/?group_id=296, Cambridge_Buckner dataset (fMRI_1) and New York dataset (fMRI_2)). fMRI_1 measures the time series resting state signal of 198 adult individuals and fMRI_2 measures 84 individuals. Both datasets were acquired using gradient-echo echo-planar-imaging on 3 Tesla scanners. For Cambridge_Buckner dataset, 119 volumes were acquired under the following parameters: TR = 3000 ms, voxel size = 3 × 3× 3 mm^3^, FOV = 216 × 216 mm^2^, 47 slices, TE = 30 ms, flip angle = 85 degrees; For New_York dataset, 192 volumes were acquired under the following parameters: TR = 2000 ms, voxel size = 3 × 3× 3 mm^3^, FOV = 192 × 192 mm^2^, 39 slices, TE = 25 ms, flip angle = 80 degrees. Only the 75 male samples were included in our analysis, to directly compare to our male transcriptome brain samples. In the replication datasets, all 198 individuals and 84 individuals were included as both male and female transcriptome samples are available. The preprocessing of the functional and anatomical images was done according to preprocessing scripts provided on the fcon_1000 website (http://www.nitrc.org/projects/fcon_1000). The preprocessing was done using FSL (Jenkinson et al., 2012) and AFNI (Cox, 2012). It consists of the following steps: 1. Realigning the functional images to correct for motion; 2. Spatially smoothing the functional images (Gaussian Kernel: FWHM 6 mm); 3. Rescaling the functional image to the mean; 4. Temporal band-pass filtering the functional image (0.01-0.08 Hz); 5. Removing linear and quadratic trends of the functional image; 6. Normalizing the functional image from subject space into Montreal Neurological Institute (MNI) 152 space based on T1 images using linear transformation; 7. Regressing out the global brain, white matter, CSF and motion parameter signal from the functional image. 8. Calculating mean signal time course for each region of interest (ROI). The masks for Brodmann ROIs were obtained from the software PickAltas (Maldjian et al., 2003) and were resampled into the same resolution as the functional images. Then frequency amplitude of low-frequency fluctuation (fALFF) was calculated as previously reported(Zou et al., 2008). Briefly, power spectrum was computed for each ROI, the square root of which is the amplitude at each frequency. The raw fALFF index was calculated as the sum amplitude within 0.01-0.08 Hz divided by the sum of amplitudes over the whole frequency range. For standardization purposes, the raw fALFF of each ROI was divided by the mean of the raw fALFF across all 10 ROIs. Finally, standardized fALFF was averaged across subjects to yield one value for each ROI. Then the same procedure was used to calculated fALFF in the 11 ROIs. Pearson correlation between the expression level of genes and the average fALFF among individuals were calculated. We repeated our main analysis by using Amplitude of Low Frequency Fluctuations (ALFF), another measure of resting state fMRI activity, and found that similar numbers of fMRI correlated genes are identified and their correlation levels are very similar with the fALFF results in either fMRI dataset and regardless of sex (Figure S7 and data not shown).

#### Replication datasets

The Brainspan RNA-seq data (www.brainspan.org) and Kang et al. microarray data (Kang et al., 2011) were used for gene expression replication. For each dataset, only samples from individuals aged five years or older were used. After this filtering, 378 samples from 11 neocortical regions in both left and right hemispheres in the Kang et al. dataset and 129 samples from 11 cortical regions from the Brainspan dataset were included.

#### Gene set enrichment analysis

Gene set enrichment analysis was performed using the PANTHER web resource to investigate the functional enrichment of genes that are positively or negatively correlated with the fMRI signal, as measured by fALFF. We used a Bonferroni correction to control the familywise error rate at 0.05 (Mi et al., 2013).

#### Identification of individual genes

To identify the individual genes with expression correlated with the fMRI signal, only the brain regions that are involved in the default mode network (DMN) were included (Buckner et al., 2008). In our RNA-seq data, 25 samples from 5 DMN regions were used (BA21, BA22, BA39, BA40 and BA9). In the Kang et al. dataset 212 samples from 6 DMN regions in both left and right hemispheres were used, and in the Brainspan dataset 72 samples from 6 DMN regions were included (IPC, OFC, MFC, ITC, STC and DFC). We considered the gene to have a reliable relationship with fALFF if the correlation exists in two of the three gene expression datasets (with two-tailed *P* < 0.005) across both of the fMRI datasets. *P* values were determined by the Fisher r-to-z transformation.

#### Identification of differentially expressed (DE) genes among human brain regions

Mapped reads from above were counted using HTseq-count (Anders et al., 2015) and the GRCh37 GTF provided in Ensembl release 67. Regional differentially expressed genes (genes that are differentially expressed between any brain region pair) were identified using both edgeR and DESeq in R/Bioconductor (False Discovery Rate of 0.05; Table S8) (Anders and Huber, 2010; Robinson et al., 2010). Final adjusted p values were combined by the Fisher’s combined probability test and 0.05 was used as final cutoff.

#### Identification of cell type specificity expression

We identified genes that preferentially are expressed in different cell types in the brain (http://web.stanford.edu/group/barres_lab/brain_rnaseq.html) (Zhang et al., 2014) Expression levels (RPKM) of the fALFF correlated genes were obtained in different cell types to investigate the preferred expression pattern of each gene. Then the cell type that contains the greatest RPKM expression was defined as the preferentially expressed cell type (Table S6). Only the fALFF-correlated genes that have expression information reported were counted and the Fisher exact test was used for the enrichment analysis of neuronal genes in the fALFF correlated genes.

#### Permutations to determine significance of enrichment of fALFF-correlated genes

To test whether the three gene expression datasets are correlated with the fMRI signals (fALFF) in general, large-scale permutation experiments were performed for each of them in turn. The procedure was as follows: We calculated the number of genes correlated with fALFF (fMRI_1 and fMRI_2) in each dataset. Next, we permuted fALFF value assignments to regions in each dataset and recalculated the correlations with fALFF (fMRI_1). We next determined the number of genes with significant correlations (p < 0.005) in the randomized dataset. We repeated the permutation and correlation calculations for fMRI_2 to obtain the final fALFF genes and overlapped the resultant lists from fMRI_1 and fMRI_2. Finally, we repeated these steps 1,000 times to obtain the distribution of the number of fALFF-correlated genes under the null model. To determine the significance of the 38 fALFF correlated genes, the same steps were used except that we coupled the overlapping brain regions with the fALFF signals among the three gene expression datasets. Specifically, BA9 is similar to dorsolateral prefrontal cortex (DFC), BA22 is similar to the superior temporal cortex (STC), and BA39 and BA40 are both considered part of the posterior inferior parietal cortex (IPC). A p-value was obtained by summing all the counts equal to or higher than the observed count of genes and then dividing that by the number of iterations.

#### Permutations to determine significance of enrichment of DE genes

To test whether differential expressed (DE) genes are more likely correlated with fALFF, permutation procedures similar to above were used except that all the DE genes were used as the background. p < 0.05 was used as significant.

#### Permutations to determine the co-expression of fALFF correlated genes

To test whether fALFF correlated genes have significantly higher co-expression levels than other genes, 10,000 permutations were performed. For each permutation, 38 genes were randomly sampled from the expressed gene list. Then the average co-expression values for each of them were calculated and compared with the average co-expression of the identified fALFF correlated genes. The p-value was obtained by summing all the counts equal to or higher than the observed count of co-expression and then dividing that by the number of iterations.

#### Permutation test for the correlation of fALFF correlated genes in the non-nervous system datasets

To test whether the fALFF correlated genes have a higher correlation with fALFF values in non-nervous system expression datasets, we used the GTEx dataset (GTEx Consortium. 2015). Specifically, gene expression from 25 non-central nervous system tissues (Adipose Tissue, Adrenal Gland, Bladder, Blood, Breast, Cervix Uteri, Colon, Esophagus, Fallopian Tube, Heart, Kidney, Liver, Lung, Muscle, Ovary, Pancreas, Prostate, Salivary Gland, Small Intestine, Spleen, Stomach, Testis, Thyroid, Uterus and Vagina) were obtained and correlations were made between gene expression in those samples and a set of fALFF values randomly drawn from the union of fMRI_1 and fMRI2. We computed where the observed Wilcoxon rank sum test U value of the correlations of these 38 genes (ranked amongst all genes in the test sets) fell in the distribution of simulated U values from 100 simulations.

#### Statistical analysis and code availability

Pearson correlations are reported (Figures 3A, B, C, D, Figures S2A, S2B, Figure S7, **Figure S8**). For Figures S4 and S6, two-sided Wilcoxon rank sum tests were performed. Fisher’s exact test was used to assess the significance of enrichment of fALFF-correlated genes in specific cell types, DE genes and the asdM12 module. Custom scripts in R were used for all analyses and are available upon request to the corresponding authors.

## ACCESSION NUMBERS

The NCBI Gene Expression Omnibus (GEO) accession number for the next-generation sequencing data reported in this paper is GSE58604.

## ACKNOWLEDGMENTS

This work was supported by a T32 to T.G.B. from NINDS grant T32NS048004; an NIMH grant (R00MH090238), a March of Dimes Basil O’Connor Starter Scholar Research Award, Once Upon a Time Foundation, CREW Dallas and a Jon Heighten Scholar in Autism Research Award at UT Southwestern to G.K.; the Yerkes National Primate Research Center (National Center for Research Resources P51RR165; currently Office of Research Infrastructure Programs/OD P51OD11132) to T.M.P.; and NIMH grants (5R37MH060233 and 5R01MH094714), an Autism Center for Excellence network grant (9R01MH100027), and the Simons Foundation (SFARI 206744) to D.H.G. Human tissue was obtained from the NICHD Brain and Tissue Bank for Developmental Disorders at the University of Maryland (NICHD contract numbers N01-HD-4-3368 and N01-HD-4-3383). The role of the NICHD Brain and Tissue Bank is to distribute tissue and therefore cannot endorse the studies performed or the interpretation of results.

### Competing financial interests

The authors declare no competing financial interests.

## Supplemental Information

**Figure S1.**
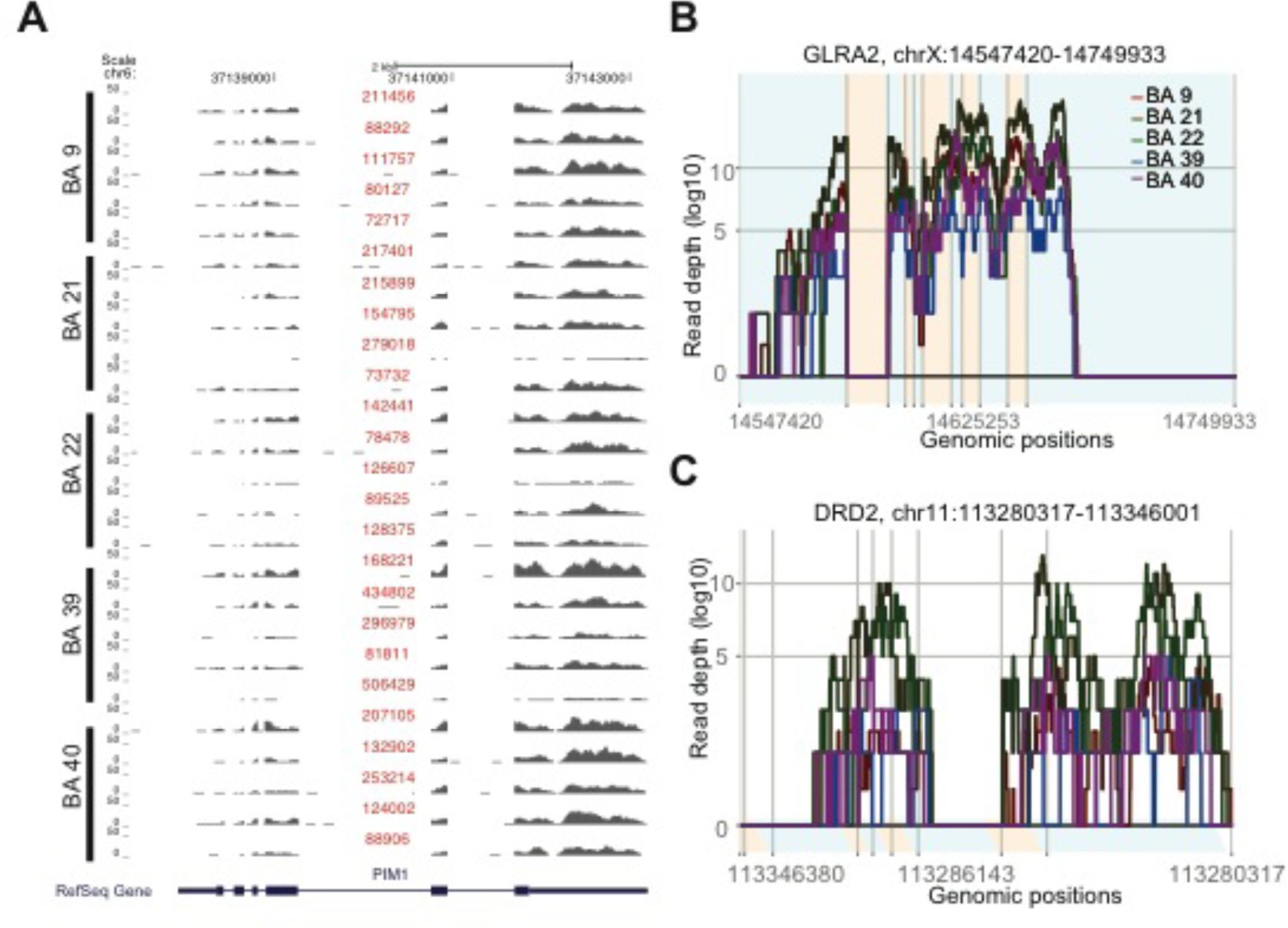
**Related to** Figure 1 Example transcriptional profiles of genes in samples from 5 brain regions (in this case 5 DMN-associated regions) in this study. (**a**) Expression of the *PIM1* gene in five cortical areas from five individuals are displayed. RPKM values for each sample are indicated in purple text above the profile and are scaled to the same height (0-400). (**b** and **c**) Example of a non-DE gene (*GLRA2*) and a DE gene (*DRD2*). Average read depth in different cortical brain regions are shown in different colors.

**Figure S2.**
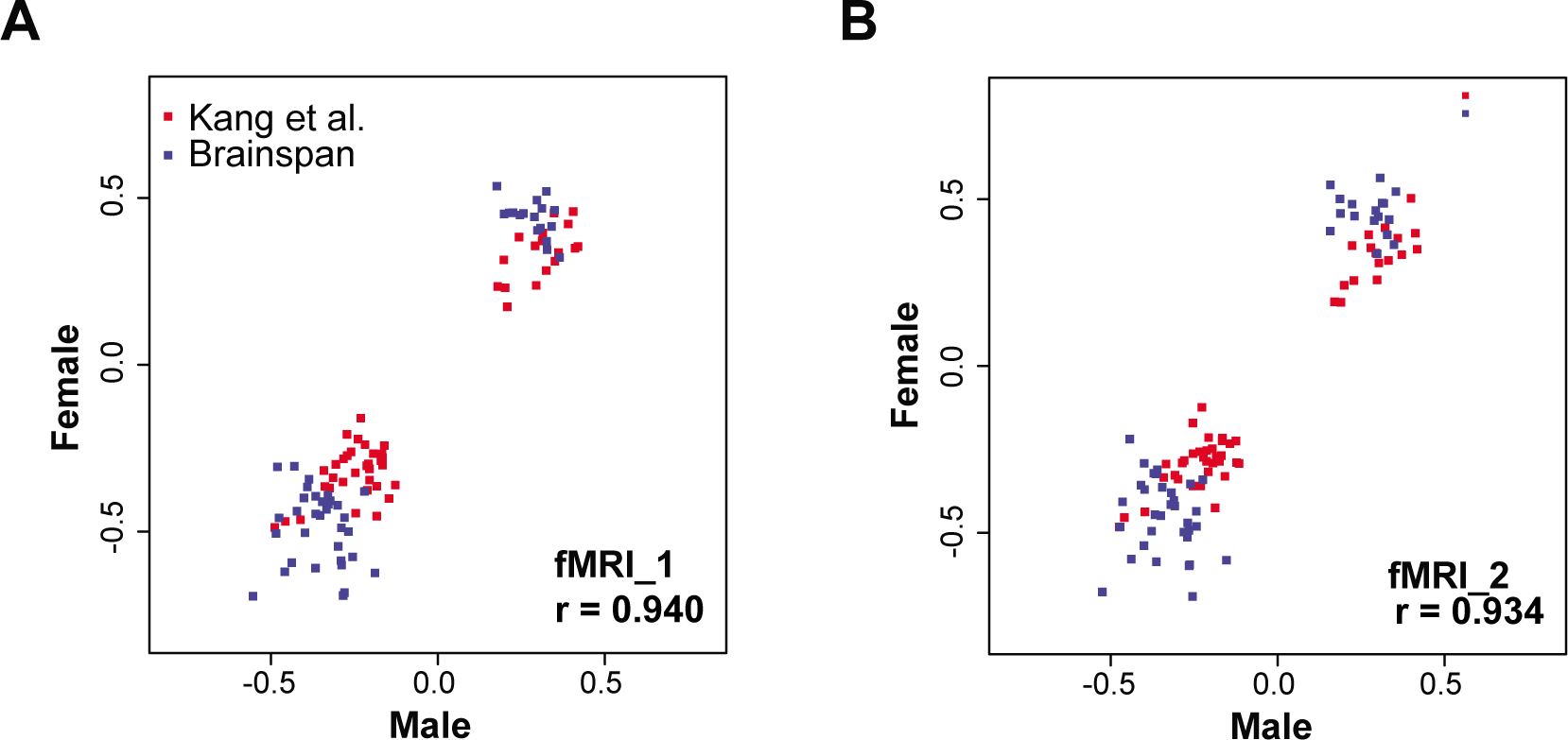
**Related to** Figure 2 Correlation between male and female samples. Genes highly correlated with fALFF in the DMN are illustrated. p < 1e-10 for all the correlations. These data demonstrate that sex is not driving the observed correlations in either (**a**) fMRI_1 dataset or (**b**) fMRI_2 dataset.

**Figure S3.**
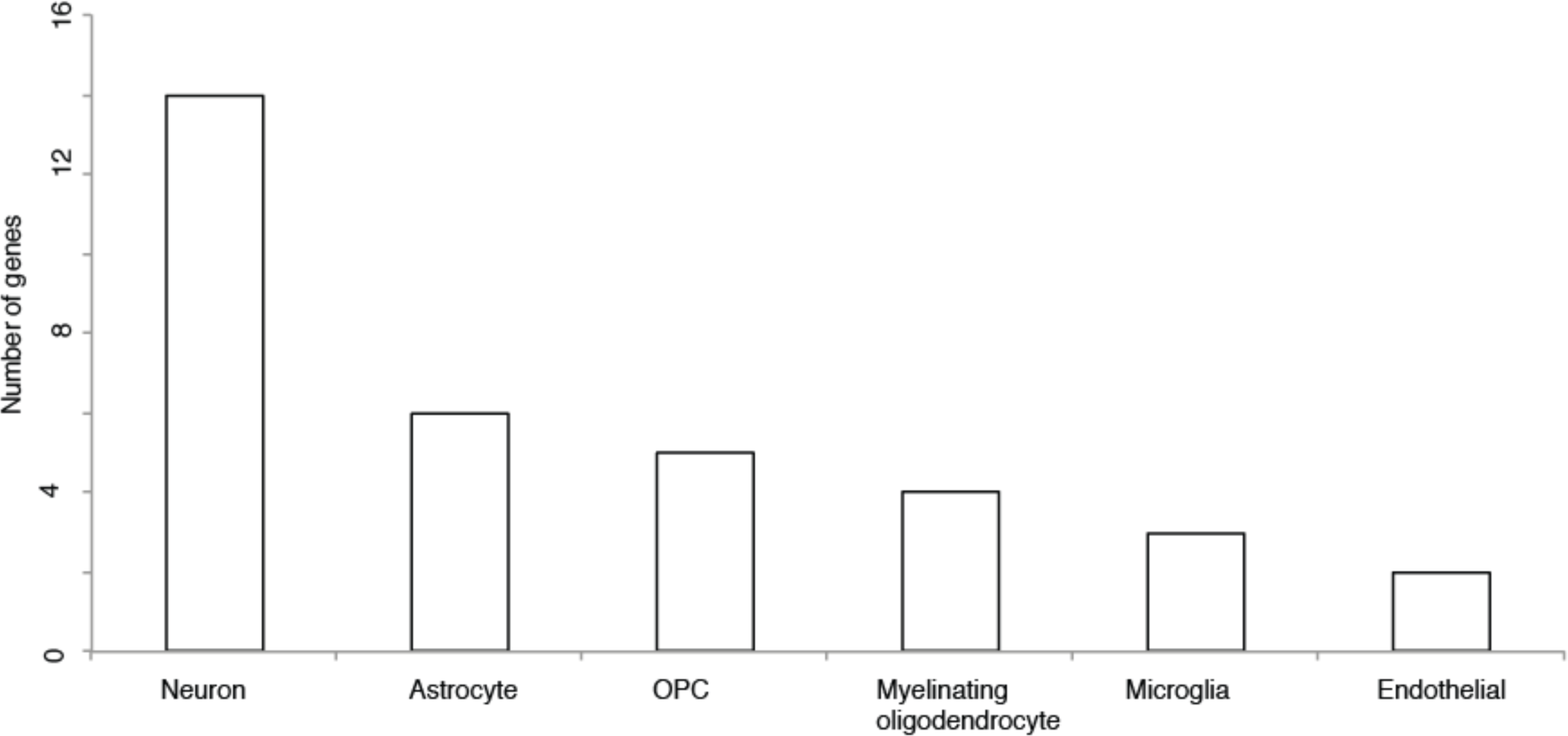
**Related to** Figure 3 Distribution of the most fALFF-correlated genes in different brain cell types in mouse. A plurality of genes is preferentially expressed in neurons. Detailed information is described in the methods and contained in Supplementary Table 6. Data were obtained as previously reported (Zhang et al., 2014).

**Figure S4.**
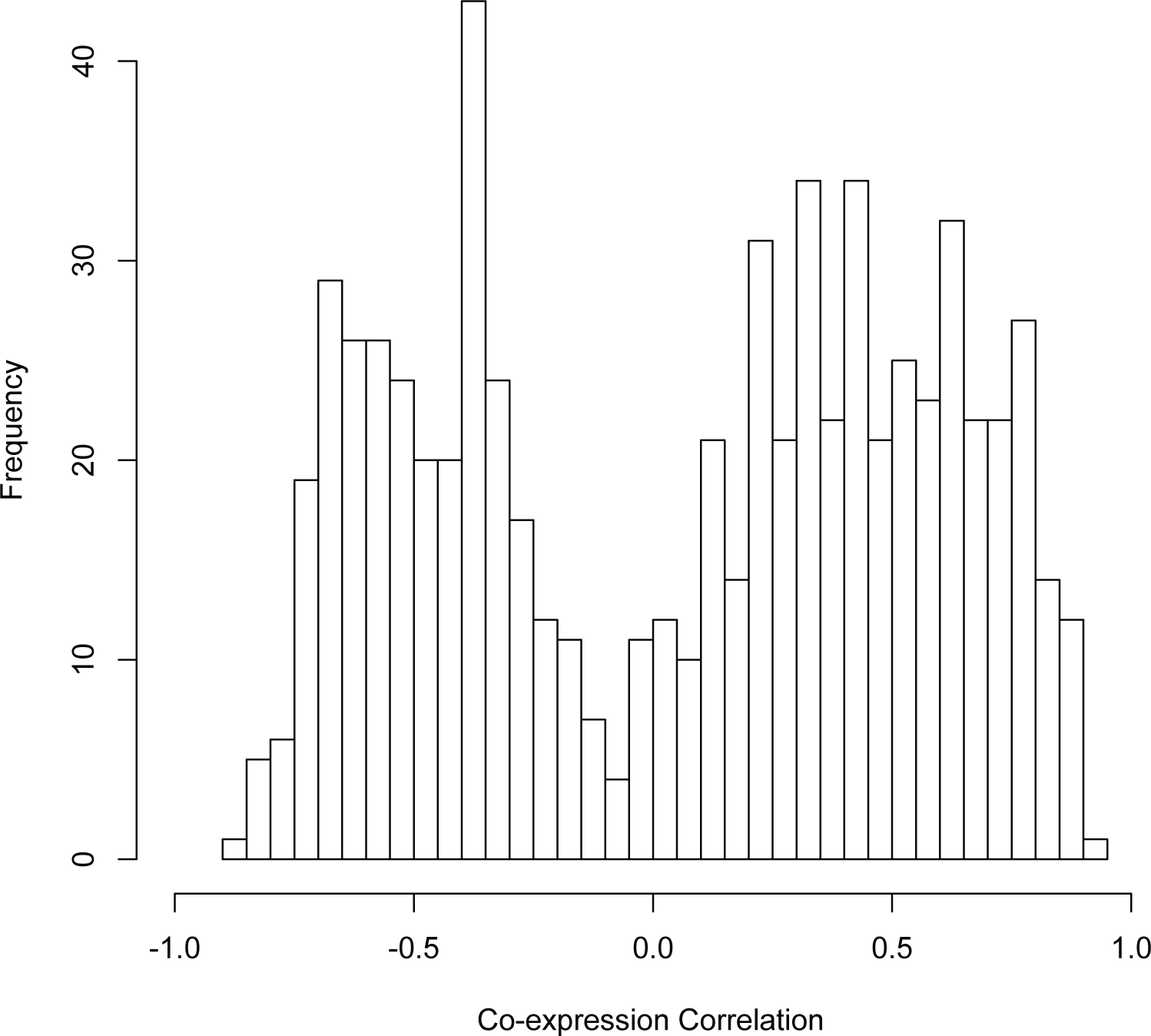
**Related to** Figure 3 Distribution of co-expression strength of the 38 fALFF-correlated genes. The co-expression among the most fALFF-correlated genes is moderately correlated in both the positive and negative directions.

**Figure S5.**
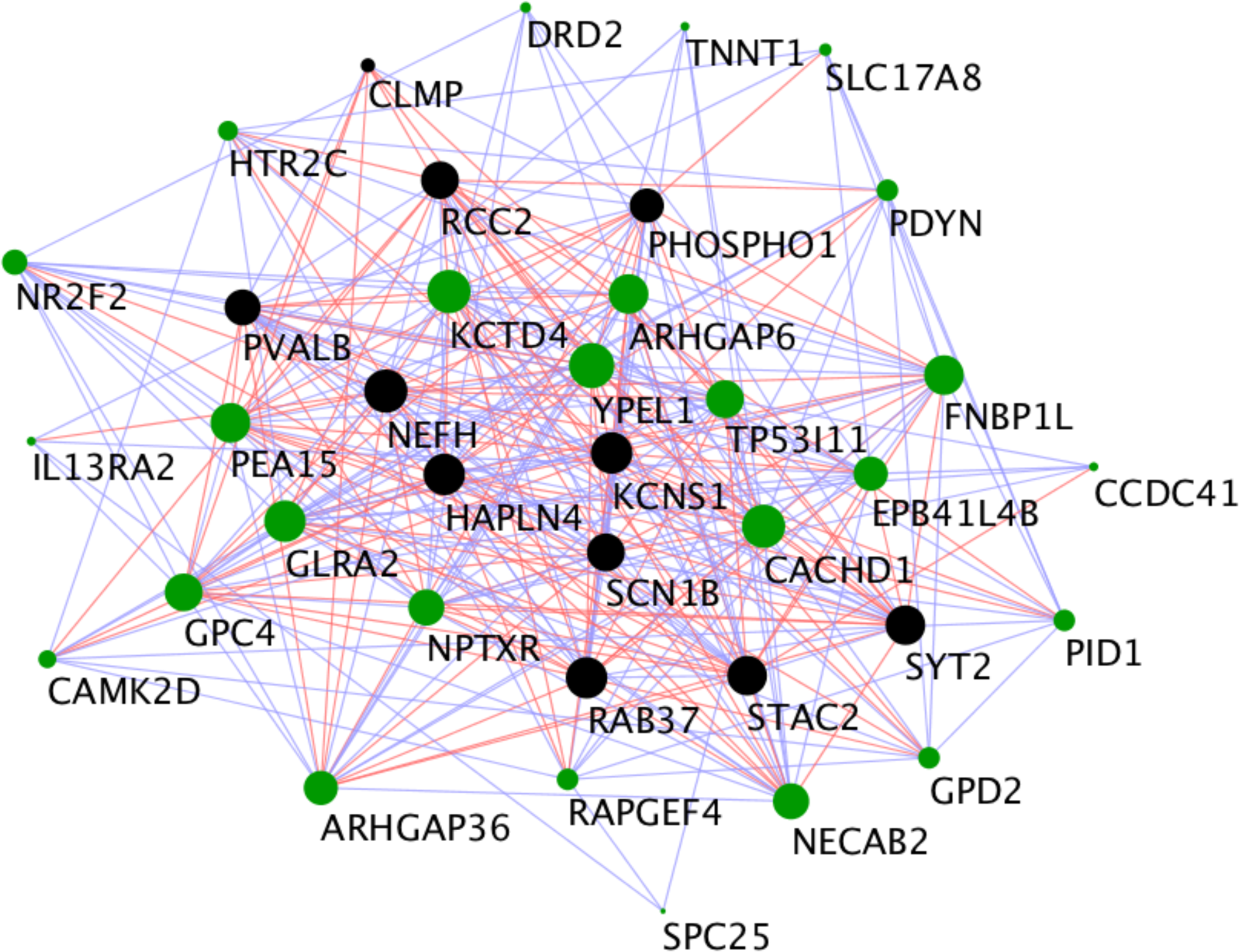
**Related to** Figure 4 fALFF-correlated genes are highly co-expressed. Blue lines indicate negative correlations (r < -0.5) and red lines indicate positive correlations (r > 0.5). The size of the points reflects the number of connected links for a particular gene. Positively fALFF-correlated genes are highlighted in black and negatively fALFF-correlated genes are highlighted in green.

**Figure S6.**
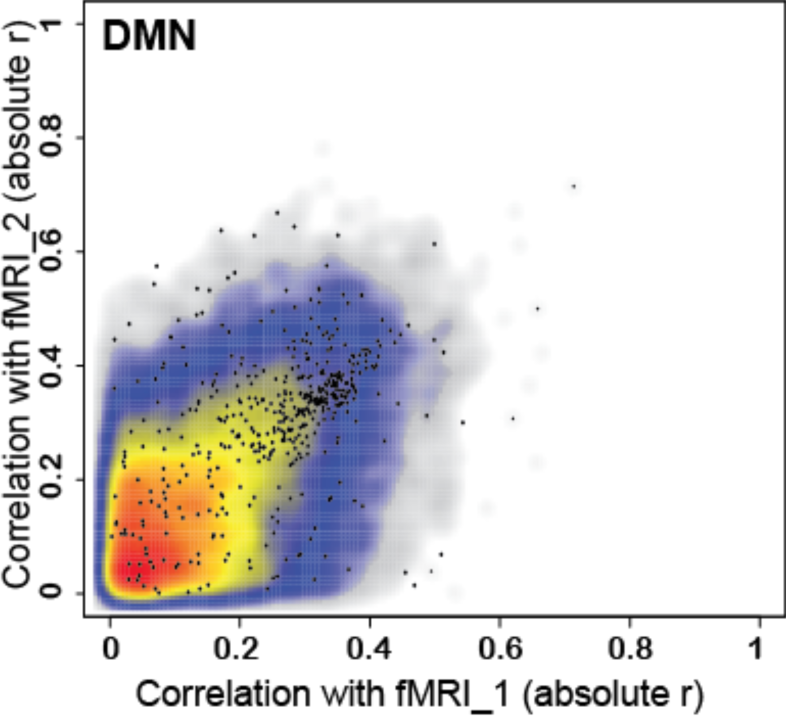
**Related to** Figure 4 Regional DE genes are more significantly correlated with fALFF. Density plots indicating the correlation of genes between the two fMRI datasets in the DMN regions. The absolute value of r is used to represent the correlation strength. Black points represent the correlation of DE genes. Other genes are plotted as a color density: grey to red indicates low to high density. p < 1e-10 in all the comparisons (Wilcoxon rank sum test).

**Figure S7.**
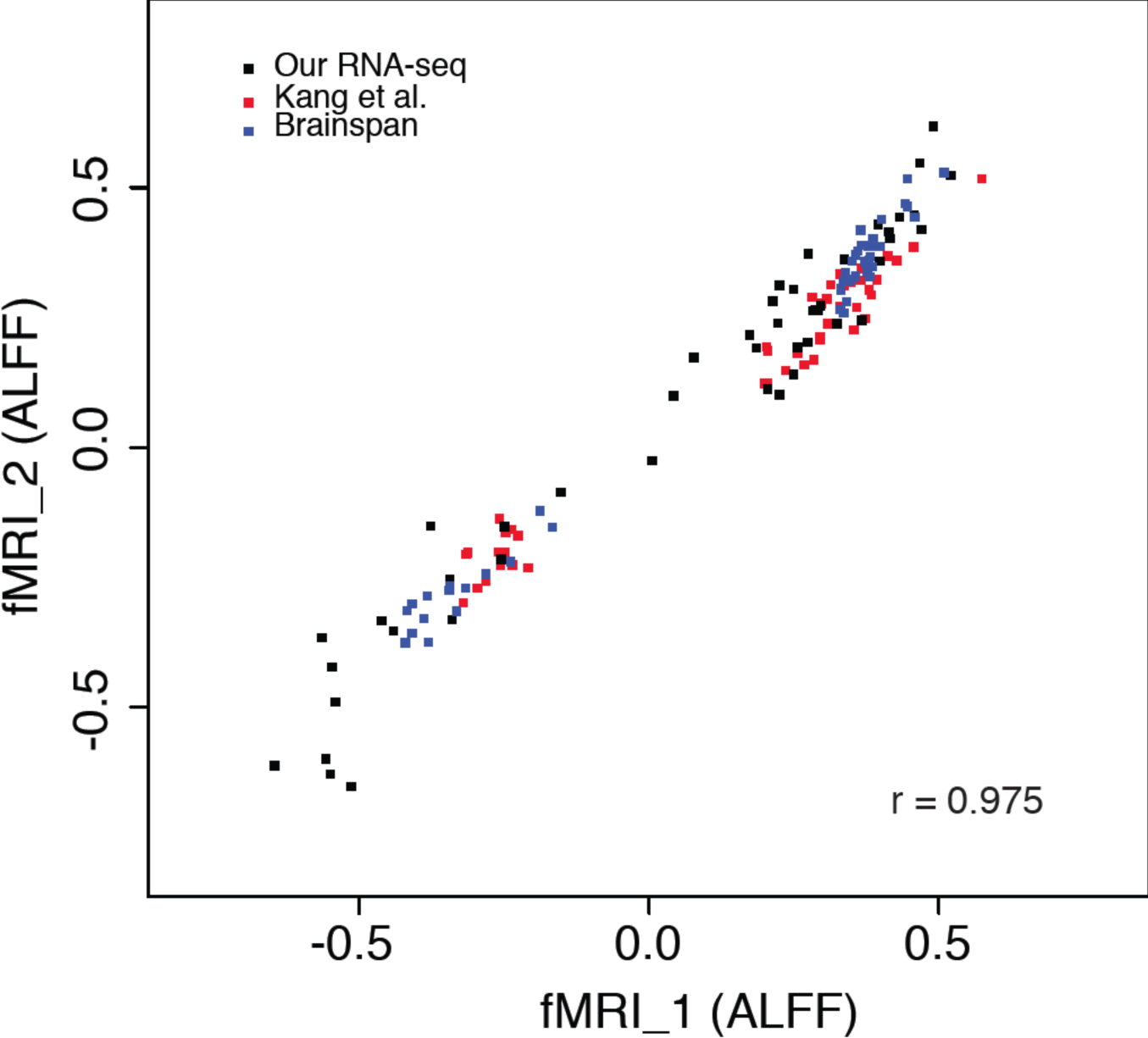
**Related to** Figure 1 Correlation between fMRI datasets using ALFF in the DMN regions. Using another measurement of brain activity (ALFF), we find the same high level of correlation between fMRI datasets (p < 1e-10).

**Table S1.**
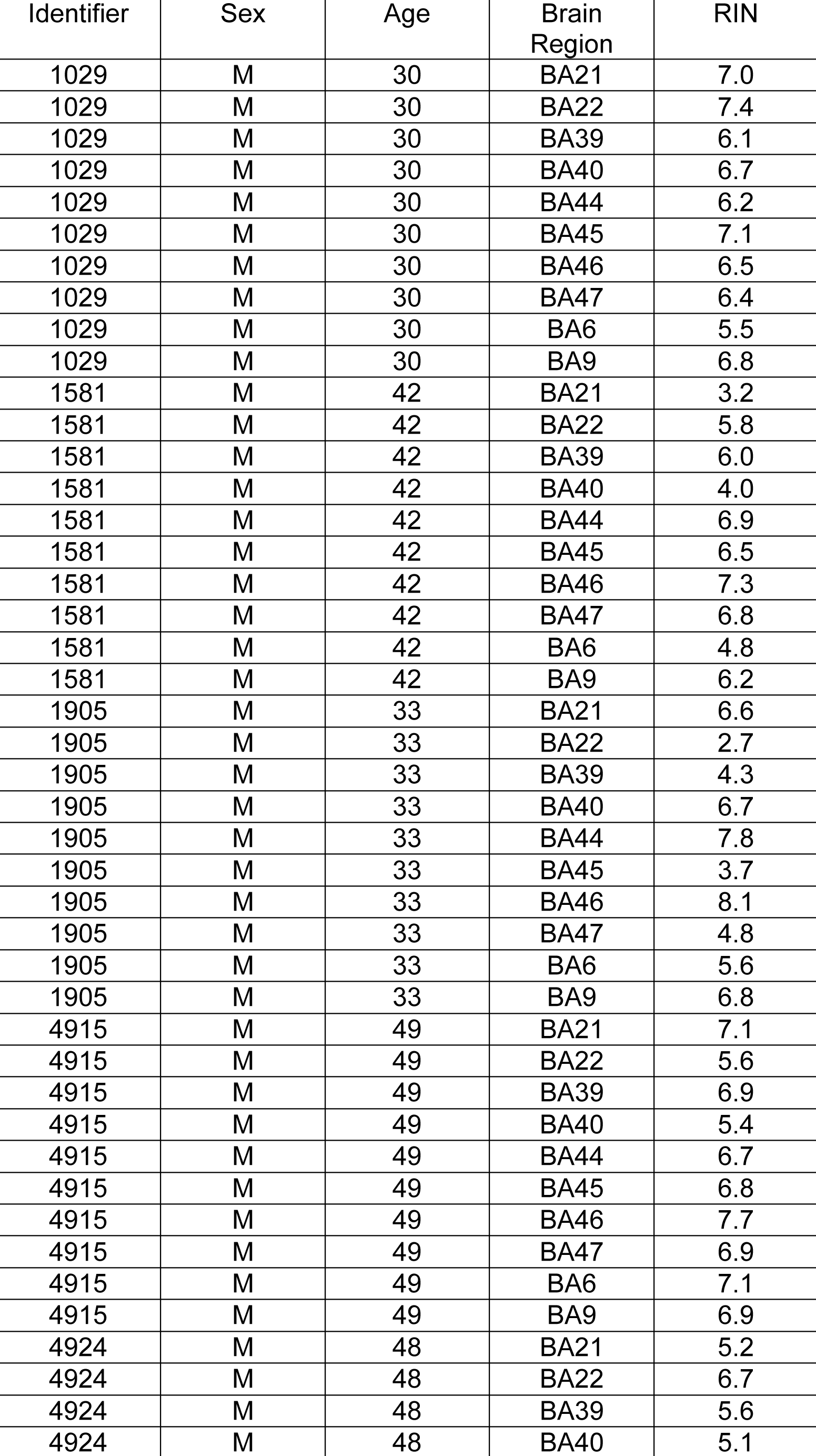
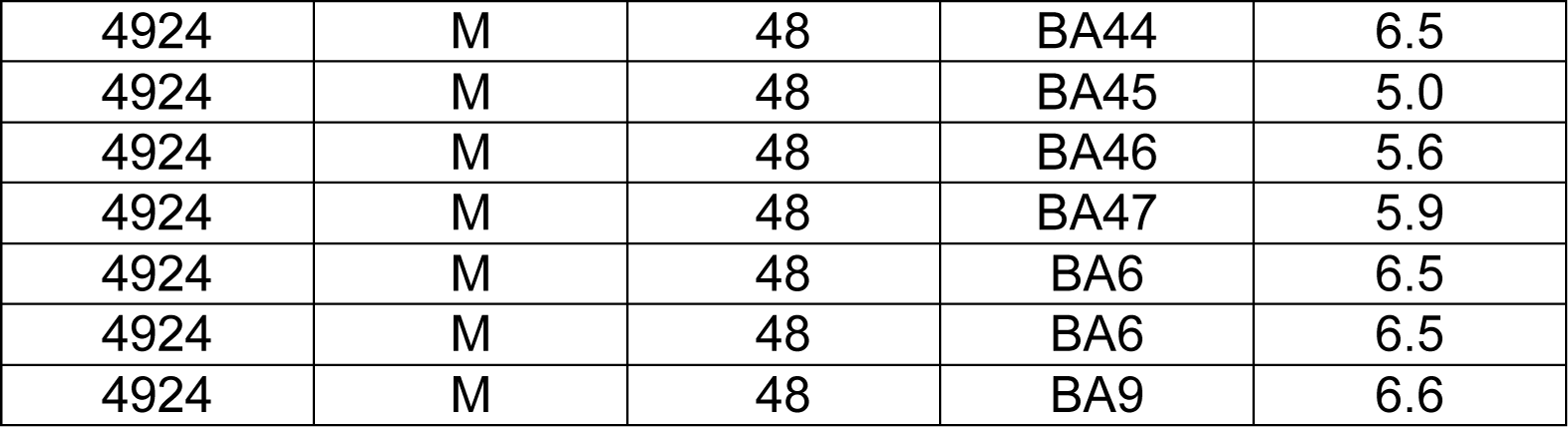
Detailed information of human neocortical samples that are used in this study.

| Identifier | Sex | Age | Brain Region | RIN |
| --- | --- | --- | --- | --- |
| 1029 | M | 30 | BA21 | 7.0 |
| 1029 | M | 30 | BA22 | 7.4 |
| 1029 | M | 30 | BA39 | 6.1 |
| 1029 | M | 30 | BA40 | 6.7 |
| 1029 | M | 30 | BA44 | 6.2 |
| 1029 | M | 30 | BA45 | 7.1 |
| 1029 | M | 30 | BA46 | 6.5 |
| 1029 | M | 30 | BA47 | 6.4 |
| 1029 | M | 30 | BA6 | 5.5 |
| 1029 | M | 30 | BA9 | 6.8 |
| 1581 | M | 42 | BA21 | 3.2 |
| 1581 | M | 42 | BA22 | 5.8 |
| 1581 | M | 42 | BA39 | 6.0 |
| 1581 | M | 42 | BA40 | 4.0 |
| 1581 | M | 42 | BA44 | 6.9 |
| 1581 | M | 42 | BA45 | 6.5 |
| 1581 | M | 42 | BA46 | 7.3 |
| 1581 | M | 42 | BA47 | 6.8 |
| 1581 | M | 42 | BA6 | 4.8 |
| 1581 | M | 42 | BA9 | 6.2 |
| 1905 | M | 33 | BA21 | 6.6 |
| 1905 | M | 33 | BA22 | 2.7 |
| 1905 | M | 33 | BA39 | 4.3 |
| 1905 | M | 33 | BA40 | 6.7 |
| 1905 | M | 33 | BA44 | 7.8 |
| 1905 | M | 33 | BA45 | 3.7 |
| 1905 | M | 33 | BA46 | 8.1 |
| 1905 | M | 33 | BA47 | 4.8 |
| 1905 | M | 33 | BA6 | 5.6 |
| 1905 | M | 33 | BA9 | 6.8 |
| 4915 | M | 49 | BA21 | 7.1 |
| 4915 | M | 49 | BA22 | 5.6 |
| 4915 | M | 49 | BA39 | 6.9 |
| 4915 | M | 49 | BA40 | 5.4 |
| 4915 | M | 49 | BA44 | 6.7 |
| 4915 | M | 49 | BA45 | 6.8 |
| 4915 | M | 49 | BA46 | 7.7 |
| 4915 | M | 49 | BA47 | 6.9 |
| 4915 | M | 49 | BA6 | 7.1 |
| 4915 | M | 49 | BA9 | 6.9 |
| 4924 | M | 48 | BA21 | 5.2 |
| 4924 | M | 48 | BA22 | 6.7 |
| 4924 | M | 48 | BA39 | 5.6 |
| 4924 | M | 48 | BA40 | 5.1 |
| 4924 | M | 48 | BA44 | 6.5 |
| 4924 | M | 48 | BA45 | 5.0 |
| 4924 | M | 48 | BA46 | 5.6 |
| 4924 | M | 48 | BA47 | 5.9 |
| 4924 | M | 48 | BA6 | 6.5 |
| 4924 | M | 48 | BA6 | 6.5 |
| 4924 | M | 48 | BA9 | 6.6 |

**Table S2.** RPKM Values. The RPKM values can be downloaded from NCBI GEO through the following reviewer link:

**Table S3.** Standardized fractional Amplitude of Low Frequency Fluctuation (fALFF) for the ten brain regions in this study.

| Brodmann area and hemisphere | fALFF (fMRI_1) | fALFF (fMRI_2) | Location | DMN | Abbreviation |
| --- | --- | --- | --- | --- | --- |
| Brodmann 21 left | 0.99 | 0.97 | Middle temporal gyrus | + | pMTG |
| Brodmann 22 left | 0.98 | 0.96 | Superior Temporal Gyrus | + | pSTG |
| Brodmann 39 left | 1.05 | 1.06 | Angular gyrus | + | AG |
| Brodmann 40 left | 1.01 | 1.05 | Supramarginal gyrus | + | SMG |
| Brodmann 44 left | 0.96 | 0.94 | Pars opercularis | – | POP |
| Brodmann 45 left | 0.99 | 1.01 | Pars triangularis | – | PTr |
| Brodmann 46 left | 1.03 | 1.04 | Middle frontal gyrus | – | MFG |
| Brodmann 47 left | 0.99 | 0.96 | Pars orbitalis | – | Porb |
| Brodmann 6 left | 0.99 | 1.01 | Pre-motor cortex | – | PMV |
| Brodmann 9 left | 1.01 | 1.01 | Dorsolateral prefrontal cortex | + | DLPFC |

**Table S4.** Standardized fractional Amplitude of Low Frequency Fluctuation (fALFF) for the brain regions in Brainspan and the Kang et al. datasets.

| Abbreviation and hemisphere | fALFF (male) | fALFF (female) | fALFF (fMRI_2, male) | fALFF (fMRI_2, female) | DMN | Description |
| --- | --- | --- | --- | --- | --- | --- |
| A1C left | 0.94 | 0.93 | 0.90 | 0.94 | – | Primary auditory (A1) cortex |
| A1C right | 0.95 | 0.95 | 0.96 | 0.95 |  |  |
| DFC left | 1.02 | 1.01 | 1.00 | 1.00 | + | Dorsolateral prefrontal cortex |
| DFC right | 1.01 | 1.02 | 0.99 | 1.00 |  |  |
| IPC left | 1.04 | 1.01 | 1.03 | 1.01 | + | Posterior inferior parietal cortex |
| IPC right | 1.06 | 1.06 | 1.09 | 1.09 |  |  |
| ITC left | 0.92 | 0.92 | 0.85 | 0.86 | + | Inferior temporal cortex |
| ITC right | 0.90 | 0.90 | 0.86 | 0.86 |  |  |
| M1C left | 1.02 | 1.04 | 1.05 | 1.02 | – | Primary motor (M1) cortex |
| M1C right | 1.05 | 1.06 | 1.10 | 1.07 |  |  |
| MFC left | 1.00 | 1.00 | 1.01 | 1.02 | + | Medial prefrontal cortex |
| MFC right | 0.99 | 0.98 | 0.97 | 0.98 |  |  |
| OFC left | 0.97 | 0.96 | 0.95 | 0.96 | + | Orbital prefrontal cortex |
| OFC right | 0.97 | 0.97 | 0.93 | 0.94 |  |  |
| S1C left | 1.03 | 1.05 | 1.11 | 1.07 | – | Primary somatosensory (S1) cortex |
| S1C right | 1.04 | 1.05 | 1.12 | 1.10 |  |  |
| STC left | 0.98 | 0.98 | 0.95 | 1.02 | + | Superior temporal cortex |
| STC right | 0.98 | 0.99 | 0.98 | 1.01 |  |  |
| V1C left | 1.06 | 1.05 | 1.04 | 1.04 | - | Primary visual (V1) cortex |
| V1C right | 1.03 | 1.04 | 1.10 | 1.03 |  |  |
| VFC left | 1.04 | 1.02 | 1.04 | 1.05 | - | Ventrolateral prefrontal cortex |
| VFC right | 1.02 | 1.01 | 0.99 | 0.99 |  |  |

**Table S5.** Detailed annotation of the 38 fALFF correlated genes. Provided as separate .xls file

**Table S6.** Cell type annotation of fALFF correlated genes. FPKM values for each cell type are indicated in the table. OPC = oligodendrocyte precursor cell; MO = myelinating oligodendroyctes; EC = endothelial cells

| Gene symbol | Astrocytes | Neuron | OPC | Newly Formed Oligo-dendrocyte | MO | Microglia | EC | Preferentially expressed cell type |
| --- | --- | --- | --- | --- | --- | --- | --- | --- |
| *CACHD1* | 19.1 | 3.9 | 4.9 | 1.6 | 1.1 | 0.1 | 11.3 | Astrocytes |
| *CAMK2D* | 16.6 | 12.4 | 6.4 | 2.1 | 0.9 | 8.5 | 5.1 | Astrocytes |
| *CCDC41* | 10.1 | 5.2 | 5.6 | 2.1 | 1.0 | 1.2 | 8.5 | Astrocytes |
| *GPC4* | 21.5 | 2.3 | 2.9 | 0.2 | 0.2 | 0.1 | 1.4 | Astrocytes |
| *HAPLN4* | 1.4 | 0.4 | 1.1 | 0.2 | 0.5 | 0.3 | 0.1 | Astrocytes |
| *PDYN* | 4.4 | 1.8 | 0.8 | 0.1 | 0.2 | 0.3 | 0.2 | Astrocytes |
| *GPD2* | 21.8 | 8.8 | 8.8 | 7.0 | 6.8 | 1.4 | 31.8 | EC |
| *TNNT1* | 0.1 | 0.1 | 0.2 | 0.1 | 0.4 | 0.3 | 0.5 | EC |
| *PIM1* | 20.0 | 10.9 | 5.4 | 1.0 | 0.9 | 80.4 | 41.7 | Microglia |
| *RCC2* | 22.4 | 30.9 | 34.6 | 32.4 | 13.0 | 65.8 | 22.5 | Microglia |
| *SCN1B* | 2.2 | 28.8 | 14.4 | 3.4 | 15.1 | 29.7 | 2.8 | Microglia |
| *PHOSPHO1* | 0.5 | 0.9 | 1.4 | 2.8 | 12.9 | 1.4 | 2.8 | MO |
| *PVALB* | 0.1 | 0.1 | 0.6 | 1.4 | 2.3 | 0.9 | 0.2 | MO |
| *RAB37* | 0.3 | 0.1 | 0.2 | 2.3 | 8.0 | 0.3 | 0.3 | MO |
| *RAPGEF4* | 1.6 | 5.9 | 5.1 | 20.3 | 59.4 | 2.4 | 25.7 | MO |
| *ARHGAP36* | 0.1 | 0.8 | 0.1 | 0.1 | 0.1 | 0.1 | 0.1 | Neuron |
| *ARHGAP6* | 0.2 | 0.8 | 0.5 | 0.1 | 0.1 | 0.1 | 0.6 | Neuron |
| *DRD2* | 0.1 | 5.8 | 0.2 | 0.1 | 0.2 | 0.1 | 0.1 | Neuron |
| *FNBP1L* | 11.5 | 54.5 | 14.1 | 2.8 | 0.2 | 1.6 | 24.1 | Neuron |
| *GLRA2* | 0.1 | 10.4 | 0.1 | 0.1 | 0.1 | 0.1 | 0.1 | Neuron |
| *HTR2C* | 0.1 | 2.0 | 0.1 | 0.1 | 0.1 | 0.1 | 0.1 | Neuron |
| *KCNS1* | 0.1 | 0.6 | 0.3 | 0.1 | 0.1 | 0.2 | 0.1 | Neuron |
| *NEFH* | 0.1 | 1.4 | 0.3 | 0.1 | 0.3 | 0.2 | 0.1 | Neuron |
| *NPTXR* | 0.6 | 43.5 | 8.1 | 1.7 | 3.9 | 3.3 | 0.3 | Neuron |
| *NR2F2* | 1.9 | 30.5 | 1.3 | 0.2 | 0.1 | 0.1 | 3.3 | Neuron |
| *SLC17A8* | 0.1 | 3.5 | 0.1 | 0.1 | 0.1 | 0.1 | 0.1 | Neuron |
| *STAC2* | 0.1 | 2.4 | 0.9 | 0.4 | 2.2 | 1.3 | 0.2 | Neuron |
| *SYT2* | 0.1 | 0.5 | 0.1 | 0.1 | 0.1 | 0.1 | 0.1 | Neuron |
| *YPEL1* | 0.9 | 11.1 | 5.1 | 2.7 | 1.9 | 1.2 | 1.4 | Neuron |
| *IL13RA2* | 0.1 | 0.1 | 0.2 | 0.1 | 0.1 | 0.1 | 0.1 | OPC |
| *KCTD4* | 1.7 | 0.9 | 5.7 | 3.0 | 1.9 | 0.1 | 0.2 | OPC |
| *NECAB2* | 0.8 | 13.9 | 24.2 | 13.8 | 2.0 | 1.0 | 0.3 | OPC |
| *PID1* | 12.7 | 9.7 | 60.2 | 12.6 | 0.4 | 5.6 | 1.9 | OPC |
| *SPC25* | 1.9 | 0.9 | 4.7 | 0.7 | 0.1 | 0.2 | 3.3 | OPC |

**Table S7.** The 9 fALFF-correlated genes that overlap with the autism gene co-expression module, asdM12.

| Ensembl ID | GeneName | Sign | Comments |
| --- | --- | --- | --- |
| ENSG00000105711 | *SCN1B* | + | Both fMRI_1 and fMRI_2 |
| ENSG00000100285 | *NEFH* | + | Both fMRI_1 and fMRI_2 |
| ENSG00000139190 | *VAMP1* | + | Only fMRI_2 |
| ENSG00000141750 | *STAC2* | + | Both fMRI_1 and fMRI_2 |
| ENSG00000177098 | *SCN4B* | + | Only fMRI_2 |
| ENSG00000172794 | *RAB37* | + | Both fMRI_1 and fMRI_2 |
| ENSG00000187664 | *HAPLN4* | + | Both fMRI_1 and fMRI_2 |
| ENSG00000100604 | *CHGA* | + | Only fMRI_2 |
| ENSG00000070159 | *PTPN3* | - | Only fMRI_2 |

**Table S8.** DE genes that are detected in the DMN regions. Provided as separate .xls file

